# Structural determination of *Streptococcus pyogenes* M1 protein interactions with human immunoglobulin G using integrative structural biology

**DOI:** 10.1101/2020.07.21.213652

**Authors:** Hamed Khakzad, Lotta Happonen, Yasaman Karami, Michael Nilges, Guy Tran Van Nhieu, Johan Malmström, Lars Malmström

## Abstract

*Streptococcus pyogenes* (Group A streptococcus; GAS) is an important human pathogen responsible for mild to severe, life-threatening infections. GAS expresses a wide range of virulence factors, including the M family proteins. The M proteins allow the bacteria to evade parts of the human immune defenses by triggering the formation of a dense coat of plasma proteins surrounding the bacteria, including IgGs. However, the molecular level details of the M1-IgG interaction have remained unclear. Here, we characterized the structure and dynamics of this interaction interface in human plasma on the surface of live bacteria using integrative structural biology, combining cross-linking mass spectrometry and molecular dynamics (MD) simulations. We show that the primary interaction is formed between the S-domain of M1 and the conserved IgG Fc-domain. In addition, we show evidence for a so far uncharacterized interaction between the A-domain and the IgG Fc-domain. Both these interactions mimic the protein G-IgG interface of group C and G streptococcus. These findings underline a conserved scavenging mechanism used by GAS surface proteins that block the IgG-receptor (FcγR) to inhibit phagocytic killing. We additionally show that we can capture Fab-bound IgGs in a complex background and identify the specific M1 epitopes targeted on live bacteria. Our results elucidate the M1-IgG interaction network involved in inhibition of phagocytosis and reveal important M1 peptides that can be further investigated as future vaccine targets.

## Introduction

GAS is a Gram-positive bacterium that causes both mild infections in the upper respiratory tract as well as severe, invasive systemic diseases including streptococcal toxic shock syndrome (STSS), necrotizing fasciitis (NF) and sepsis [1]. Worldwide, GAS is responsible for an estimated 700 million cases each year, of which 650,000 progress to severe invasive infections with an associated mortality of 25% [2], making GAS one of the most predominant bacterial pathogens to humans. To cause invasive infections, GAS expresses a wide range of virulence factors to evade human defense mechanisms. These virulence factors mostly consist of secreted or surface-associated proteins that target proteins and protein complexes of the innate and adaptive immune systems [3, 4, 5].

Specifically, GAS expresses several IgG degrading enzymes [6, 7, 8] and Fc-binding proteins [9, 10, 11] that target immunoglobulins G (IgGs), key players of the humoral immune response. In human plasma, IgG is the most abundant immunoglobulin isotype. The four subclasses IgG1-4 are highly conserved in sequence and structure, with significant non-random differences, especially in IgG3 with an extended hinge-region [12]. All subclasses are composed of two conserved heavy and light chains connected by a varying number of disulfide bonds. The heavy chain is composed of three constant domains (CH3, CH2, CH1), and one variable domain (VH) close to the N-terminus. The light chain is composed of one variable (VL) and one constant domain (CL). The CH2 and CH3 domains form the fragment crystallizable (Fc)-domain connected to CH1 via the hinge-region. Moreover, the CH1 and VH, together with the light chain VL domain, constitute the fragment antigen-binding (Fab)-domains (**Figure 1**).

**Figure 1:**
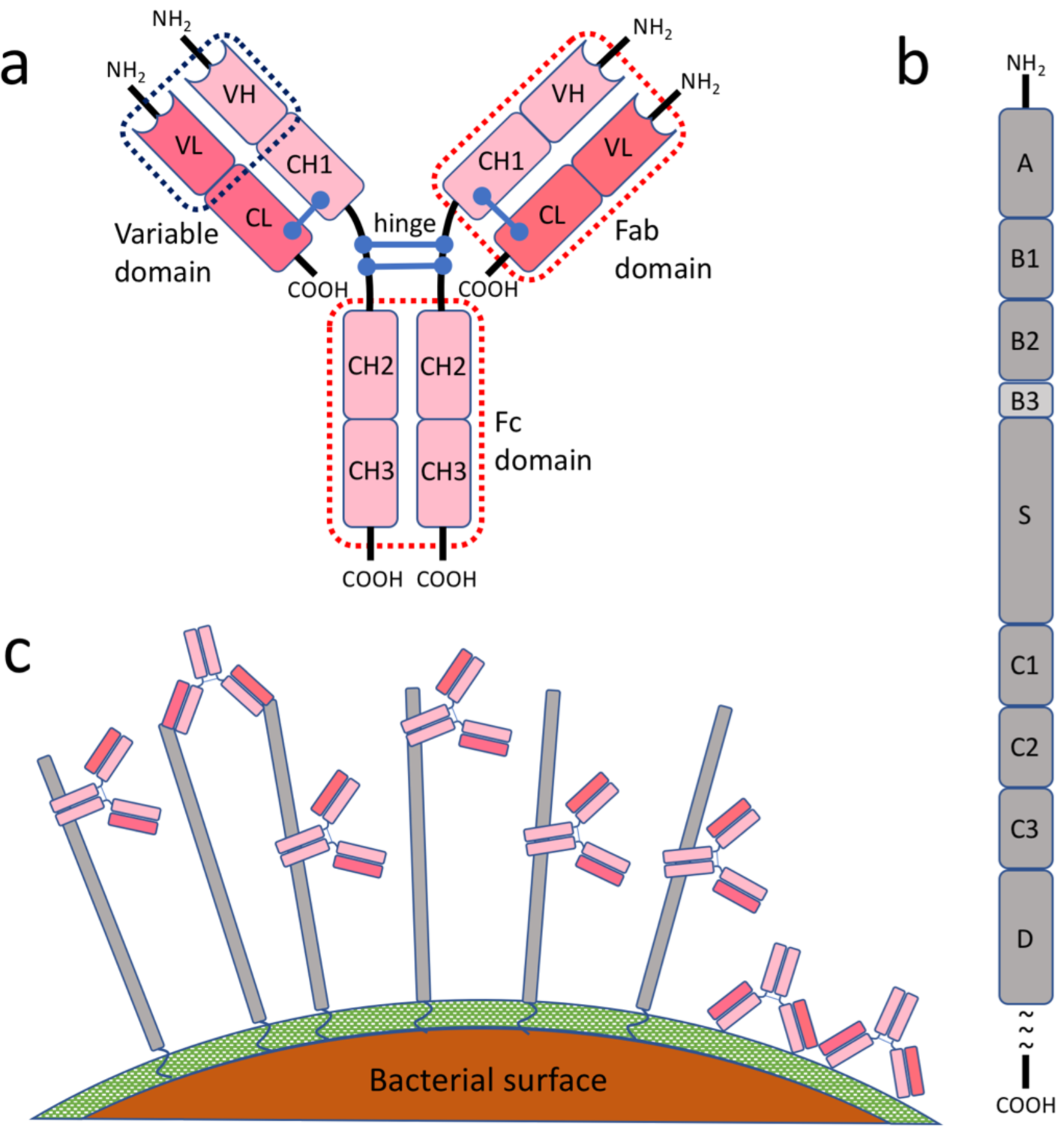
The schematic view of the studied system. **(a)** The general structure of human IgG. **(b)** Important domains of the M1 protein, including the hypervariable domain (A), fibrinogen binding domains (B1/B2), S-region, and albumin-binding domains (C1/C2/C3). **(c)** IgG-orientations and -interactions with the M1-protein on the surface of GAS. Three major interactions are shown, including Fc- and Fab-mediated interactions, as well as opsonizing antibodies bound to the surface of the bacteria *[16]*. M1 is shown in gray while IgGs are in pink. The peptidoglycan layer on the surface of bacteria is shown in green.

Together with IgGs, the complement proteins of the innate immune system have a crucial role in combatting pathogens. During the classical pathway of the complement system activation, blood plasma IgGs bind to bacterial antigens via the VL and VH of the Fab-domains. The respective Fc-domains are recognized by the complement C1q, which in turn binds to Fc-receptors on phagocytic cells initializing phagocytosis. To prevent IgG-mediated opsonization and phagocytosis, GAS uses two main classes of virulence factors: the M proteins and the IgG-degrading enzymes IdeS and EndoS/EndoS2 [7, 13]. In the family of M proteins, antigenic variation has resulted in more than 200 serotypes, but only a few are frequently associated with invasive disease, with M1 being the most prevalent serotype [14]. Moreover, to physically protect its vulnerable antigenic epitopes, GAS scavenges human plasma proteins to form a surrounding coat-like barrier [4, 15, 13], and among these interactions specifically binds IgG-molecules via their Fc-domains rendering these inaccessible for Fc-receptors [16].

Targeted cross-linking mass spectrometry (TX-MS) relies on computational structural models to score sets of targeted cross-linked peptide signals acquired using a combination of mass spectrometry acquisition techniques such as data-dependent acquisition (DDA), data-independent acquisition (DIA) and high-resolution MS1 (hrMS1). We have previously demonstrated the utility of TX-MS by creating a high-resolution quaternary model of a 1.8 MDa protein complex composed of the M1 protein and ten human plasma proteins [15]. Among these interactions, the M1-IgG binding is found to be environment-specific, binding via Fab-domains under antibody-rich conditions such as plasma, or via Fc-domains in an antibody-poor environment such as saliva [16]. We have previously shown that all human IgG subclasses can bind to the M1 protein [4]; however, the molecular details of these interactions are not fully understood.

Here, we determined the interaction of the M1 protein with human IgGs in its native environment on the bacterial surface by cross-linking the intact live bacteria in human plasma. We characterized this sophisticated mechanism in an unfractionated, complex sample using an integrative structural biology approach combining TX-MS and 2 **μ**s MD simulations. While TX-MS helped to discover the binding sites in either protein, MD simulations revealed the accurate interaction networks and the strength of the interaction.

## Methods

### Computational modeling

The UniProt accession numbers for the M1 protein and the IgG subclasses 1-4 are Q99XV0, P01857, P01859, P01860, and P01861, respectively. Using Rosetta Software suit [17], comparative models have been generated for human IgG subclasses based on crystal structures deposited in protein data bank with PDB IDs 1hzh, 4haf, 5w38, and 4c54 for IgG1, IgG2, IgG3, and IgG4, respectively. For IgG1, we modeled the gap in the hinge region using DaReUS-Loop [18, 19], re-packed the sidechains of the full structure, and made the final set of models with rosetta relax protocol considering the disulfide bridges as the input constraints.

For computational modeling of GAS M1 protein, we first separated the coiled-coil domain from signal peptide and anchor, then searched them through HHpred [20] to find homologous structures. Then, we used the Rosetta comparative modeling protocol (RosettaCM [21]) to model each domain from its homologs separately and produced 10K models per domain. To filter out the models, we used the Rosetta energy function [22, 17]. The final model is generated by another comparative modeling run, generating 10K models of the full-length protein filtered out using the Rosetta score. The length of the final model was 489.5 Å, which was aligned with our observation based on electron microscopy images [23]. To our knowledge, this is the first accurate full-length model of M1 protein.

To analyze the interaction of M1 and IgG1-4, we used TX-MS, which combines all three MS acquisition data (hrMS1, DDA, and DIA) with computational models. Accordingly, a machine learning algorithm based on hrMS1 data and a training set containing isotopic patterns of previously known XLs is used to detect potential binding interfaces. Several docking models were then generated for each binding interface, using the RosettaDock protocol [24]. These models were filtered out using cross-linking data from DDA samples, and models that were highly supported by XLs have been selected. Finally, all computational XLs below a cut-off threshold of 32 Å were produced for selected models and analyzed by DIA data.

### Cross-linking of plasma adsorption samples

For cross-linking, pooled normal human plasma was adsorbed onto the surface of *S. pyogenes* bacteria, as described [4, 15]. The *S. pyogenes* M1 serotype strain SF370 from the American Type Culture Collection (ATCC; strain reference 700294), was grown at 37 °C, 5% CO_2_ to mid-exponential phase (OD_620nm_ ∼ 0.4) in TH broth supplemented with 0.3% (w/v) yeast extract. The cells were harvested by centrifugation (1900×g, 10 min, 22°C), washed with phosphate-buffered saline (PBS, 10 mM phosphate buffer, 2.7 mM potassium chloride, 137 mM sodium chloride (Sigma)), recentrifuged (1900×g, 5 min, 22°C), and resuspended to an approximate concentration of 1 × 10^9^ colony forming units ml^−1^. Four hundred microliters of pooled normal human plasma from healthy donors (Innovative Research) supplemented with a final concentration of 10 μM argatroban (Sigma) was mixed with 100 μl of bacteria and incubated at 37 °C 30 min 500 rpm.

The bacteria with adsorbed plasma proteins were harvested by centrifugation (1900 × g, 5 min, 22°C) and washed twice with PBS and finally resuspended in 200 μl of PBS for cross-linking. Heavy/light disuccinimidyl-suberate cross-linker (DSS-H12/D12, Creative Molecules Inc.) resuspended in 100% dimethylformamide (Sigma) was added to final concentrations of 0.25, 0.5, 1.0 and 2.0 mM in duplicates, and incubated for 1 h, 37 °C, 900 rpm. The cross-linking reaction was quenched with a final concentration of 50 mM ammonium bicarbonate (Sigma) at 37 °C, 15 min, 900 rpm. The bacterial surface proteins with attached plasma proteins were released by limited proteolysis with 2 μG trypsin (Promega) /37 °C, 1 h, 800 rpm) prior to cell debris removal by centrifugation (1900×g, 15 min) and subsequent supernatant recovery. Two hundred microliters of the supernatant were recovered, and any remaining bacteria were killed by heat inactivation (85 °C, 5 min) prior to sample preparation for mass spectrometry.

### Sample preparation for mass spectrometry

The sample preparation for mass spectrometry was done as described [4, 15]. Briefly, the samples were denatured in 8 M urea-100 mM ammonium bicarbonate (both Sigma), and the cysteine bonds reduced with 5 mM tris(2-carboxyethyl)phosphine (Sigma) (37 °C, 30 min). The cysteines were alkylated with 5 mM iodoacetamide (Sigma) (22 °C, 60 min), and the samples subsequently digested using sequencing-grade lysyl endopeptidase (37 °C, 2 h) (Wako). The samples were diluted with 100 mM ammonium bicarbonate to a final urea concentration of 1.5 M and followed by digestion using trypsin (Promega) (37 °C, 18 h). Digested samples were acidified with 10% formic acid to a pH of 3.0, and the peptides were subsequently purified with C18 reverse-phase spin columns according to the manufacturer’s instructions (Macrospin columns, Harvard Apparatus). Dried peptides were reconstituted in 2% acetonitrile and 0.2% formic acid prior to MS analyses.

### MS experiments

All peptide analyses were performed on a Q Exactive HFX mass spectrometer (Thermo Scientific) connected to an EASY-nLC 1200 ultra-high-performance liquid chromatography system (Thermo Scientific). For data-dependent analysis (DDA), peptides were separated on an EASY-Spray column (Thermo Scientific; ID 75 μm× 25 cm, column temperature 45 °C) operated at a constant pressure of 800 bar. A linear gradient from 4% to 45% of 0.1% formic acid in 80% acetonitrile was run for 50 min at a flow rate of 300 nl min^−1^. One full MS scan (resolution 60,000@200 m/z; mass range 350 to 1600 m/z) was followed by MS/MS scans (resolution 15,000@200 m/z) of the 15 most abundant ion signals. The precursor ions were isolated with 2 m/z isolation width and fragmented using higher-energy collisional-induced dissociation at a normalized collision energy of 30. Charge state screening was enabled, and precursors with an unknown charge state and singly charged ions were excluded. The dynamic exclusion window was set to 15 s and limited to 300 entries. The automatic gain control was set to 3e6 for MS and 1e5 for MS/MS with ion accumulation times of 110 and 60 ms, respectively. The intensity threshold for precursor ion selection was set to 1.7e4.

For data-independent acquisition (DIA), peptides were separated using an EASY-Spray column (Thermo Scientific; ID 75 μm× 25 cm, column temperature 45 °C) operated at a constant pressure of 800 bar. A linear gradient from 4% to 45% of 0.1% formic acid in 80% acetonitrile was run for 110 min at a flow rate of 300 nl min^−1^. A full MS scan (resolution 60,000@200 m/z; mass range from 390 to 1210m/z) was followed by 32 MS/MS full fragmentation scans (resolution 35,000@200 m/z) using an isolation window of 26 m/z (including 0.5 m/z overlap between the previous and next window). The precursor ions within each isolation window were fragmented using higher-energy collisional-induced dissociation at a normalized collision energy of 30. The automatic gain control was set to 3e6 for MS and 1e6 for MS/MS with ion accumulation times of 100 ms and 120 ms, respectively.

For high-resolution MS1 (hrMS1), peptides were separated using an EASY-Spray column (Thermo Scientific; ID 75 μm× 25 cm, column temperature 45 °C) operated at a constant pressure of 800 bar. A linear gradient from 4% to 45% of 0.1% formic acid in 80% acetonitrile was run for 60 min at a flow rate of 300 nl min^−1^. High-resolution MS scans (resolution 240,000@200 m/z; mass range from 400 to 2000m/z) were acquired using automatic gain control set to 3e6 and a fill time of 500 ms.

### MD simulations

We performed MD simulations on the two complexes of M1(S-domain peptide)-IgG1(Fc) and M1(A-domain peptide)-IgG1(Fc). The starting conformations correspond to two different binding sites for the Fc domain of IgG1 (residues T108-K330) on the protein M1: (*i*) residues L108-K123 of M1(A-domain) with peptide sequence: LETKLKELQQDYDLAK, and (*ii*) residues G208-Q223 of M1(S-domain) with peptide sequence: GNAKLELDQLSSEKEQ. MD simulations were carried out with the Gromacs 2018.3 [25] using the amber99sb forcefield parameter set: (*i*) hydrogen atoms were added, (*ii*) Na^+^ and Cl^-^ counter-ions were added to reproduce physiological salt concentration (150 mM solution of sodium chloride), and (*iii*) the solute was hydrated with a triclinic box of explicit TIP3P water molecules with a buffering distance of up to 12Å. Missing residues were added using MODELLER 9.24 [26].

First, we performed 5000 steps of minimization using the steepest descent method keeping only protein backbone atoms fixed to allow protein side chains to relax. After that, the system was equilibrated for 300 ps at constant volume (NVT) and for further 1 ns using a Langevin piston (NPT) [27] to maintain the pressure, while the restraints were gradually released. For every complex, two replicates of 500 ns, with different initial velocities, were performed in the NPT ensemble. The temperature was kept at 310 K and pressure at 1 bar using the Parrinello-Rahman barostat with an integration time step of 2.0 fs. The Particle Mesh Ewald method (PME) [28] was employed to treat long-range electrostatics, and the coordinates of the system were written every 100 ps. The root mean square deviations (RMSD) of the studied complexes were measured along simulation time for all the replicates (***Supplementary Figure 4***). All systems were fully relaxed after 100 ns. Consequently, the last 400 ns of each replicate were retained for subsequent analyses. Moreover, we calculated the residue root mean square fluctuations (RMSF) over the last 400 ns of every simulation (***Supplementary Figure 5***).

### COMMA2 analysis

For every studied system, COMMA2 was applied to the last 400 ns of the two replicates of MD simulations, and communication blocks were extracted. COMMA2 identifies pathway-based communication blocks (CBs^path^), *i.e*., groups of residues that move together, and are linked by non-covalent interactions, and clique-based communication blocks (CBs^clique^), *i.e.*, groups of residues that display high concerted atomic fluctuations, and that are close in 3D space (see [29, 30] for formal definitions and detailed descriptions).

## Results

### TX-MS structural analysis

Here, we elucidate the complex network of M1-IgG interactions *ex vivo* arising in pooled normal human plasma adsorbed on the surface of live bacteria, mimicking a scenario during invasive infections. **Figure 1** shows a schematic representation of the applied approach. To study the interaction of dimeric, coiled-coil multidomain (domains A-D, **Figure 1-b**) M1 protein with human IgGs, we incubated live bacteria in human plasma, allowing for Fab- and Fc-mediated binding of the plasma IgG-molecules to the M1 protein or the opsonic, Fab-mediated binding to the surface of the bacteria (**Figure 1-c**). A mutant strain (ΔM1) lacking the M1 protein was used as a negative control (***Supplementary Figure 1***). The formed, native interactions were captured by chemical cross-linking followed by mass spectrometric identification of cross-linked peptides. Here, we produced comparative models for the M1 protein and the full-length IgG1, using RosettaCM protocol, as explained in the Methods section.

An initial DIA analysis of pooled human plasma proteins absorbed to the streptococcal surface revealed that the four subclasses of IgG are highly abundant with different intensities in the MS samples (***Supplementary Figure 1***). The measured abundance distribution deviates from the distribution in human plasma (IgG1 > IgG2 > IgG3 > IgG4), confirming that the subclasses IgG1 and IgG3 are the major targets of *S. pyogenes*, due to a combination of Fab- and Fc-bound IgGs. Moreover, data analysis of DDA samples using the sequence of the M1 protein and the heavy chains of all IgG subclasses resulted in the identification of several inter- and intra-cross-links (XLs). Accordingly, we obtained 21 distinct XLs, of which 17 are inter XLs (between the M1 protein and the different IgG subclasses), which were considered for modeling refinement. The majority of inter XLs (10 out of 17) support the Fc-mediated binding of IgGs to M1, while the rest indicate novel Fab-mediated interactions. The presence of specific M1 IgG-molecules in commercial pooled normal human plasma has been described [4]; however, the specific epitopes they target has remained unknown.

To elucidate the Fc-binding interface, we used TX-MS based on hrMS1 and DDA data that we combined with previously determined crystal structures of human IgGs and the partial peptides of M1 (A- and S-domains) [15] (see Methods). As demonstrated in **Figure 2**, all IgG subclasses bind to the M1 protein through a specific site in the IgG CH3 domain, which is essential for binding to FcγR receptors. This IgG-binding site has previously been shown to bind to a variety of proteins other than Fc-receptors, including the streptococcal protein G [31, 32] and the staphylococcal protein A [33], both of which are commercially available as tools to capture IgG-molecules via their Fc-domains. **Figure 3** shows the binding-site comparison between the M1-IgG1 model and the C2 fragment of the streptococcal protein G on the IgG Fc-domain. Accordingly, both M1 and protein G share the same binding site on the IgG Fc-domain and bind to all IgG subclasses [32].

**Figure 2:**
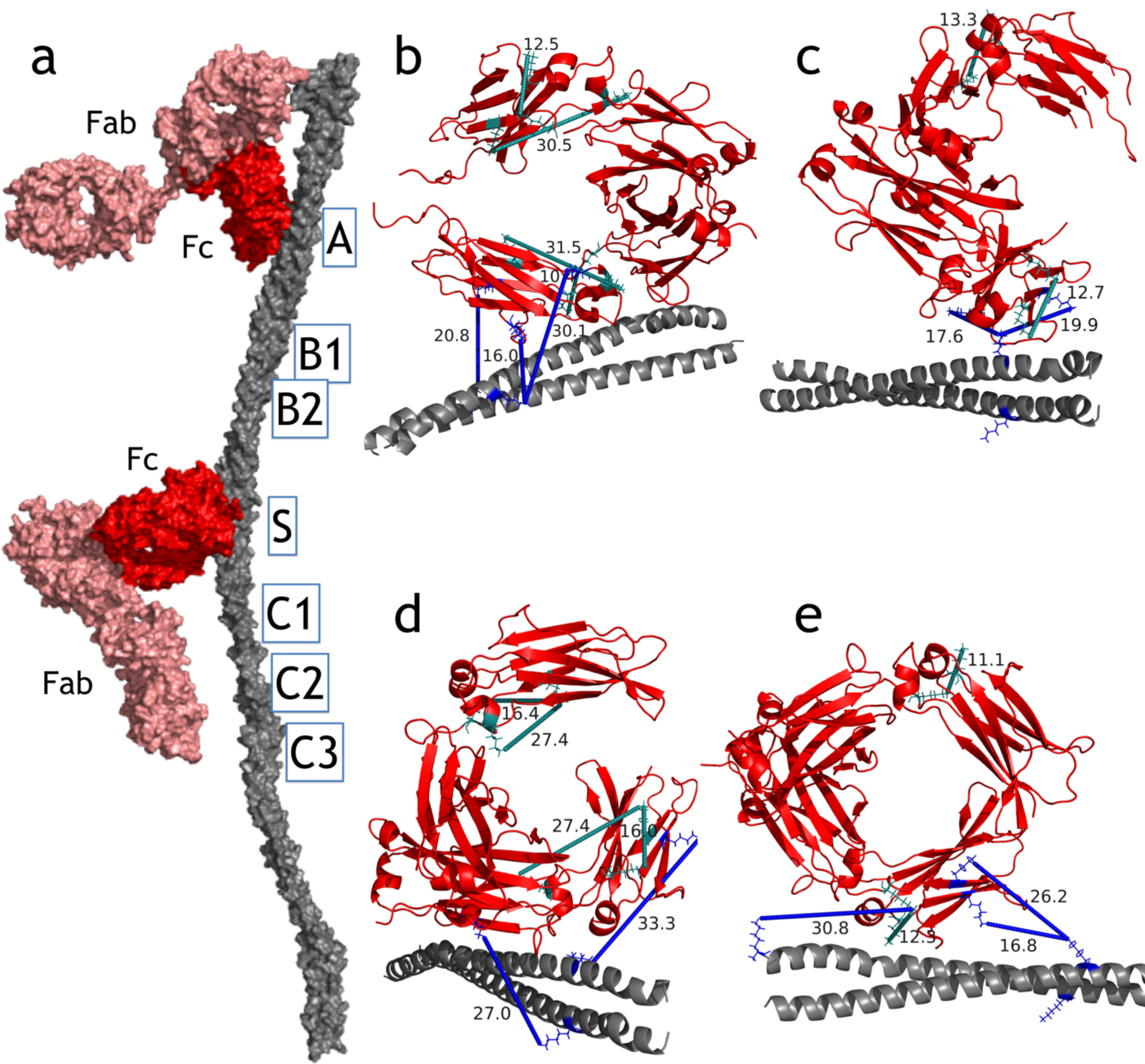
M1 interactions with IgG subclasses. **(a)** The model for M1 (A- and S-domains)-IgG1 full length non-immune Fc-mediated interaction. Important domains of the M1 protein are shown, where A- and S-domains play a crucial role. The molecular details of the Fc-binding site are shown in panel **(b)** where inter- and intra-XLs are mapped on the protein complex. **(b-e)** The S-domain of the M1-protein interacts with the Fc-region of IgG1-4. All models are generated by the TX-MS workflow based on cross-link constraints derived from MS data. Intra-XLs for each IgG are shown in cyan, while inter-XLs between the two proteins are in blue. The binding interfaces of the M1 protein on the Fc-domain of all IgG subclasses are similar and mediated via the CH2 and CH3 domains. This region is involved in binding to IgG Fc-receptors (FcγR), predicted to be inhibited by the M1 protein interaction.

**Figure 3:**
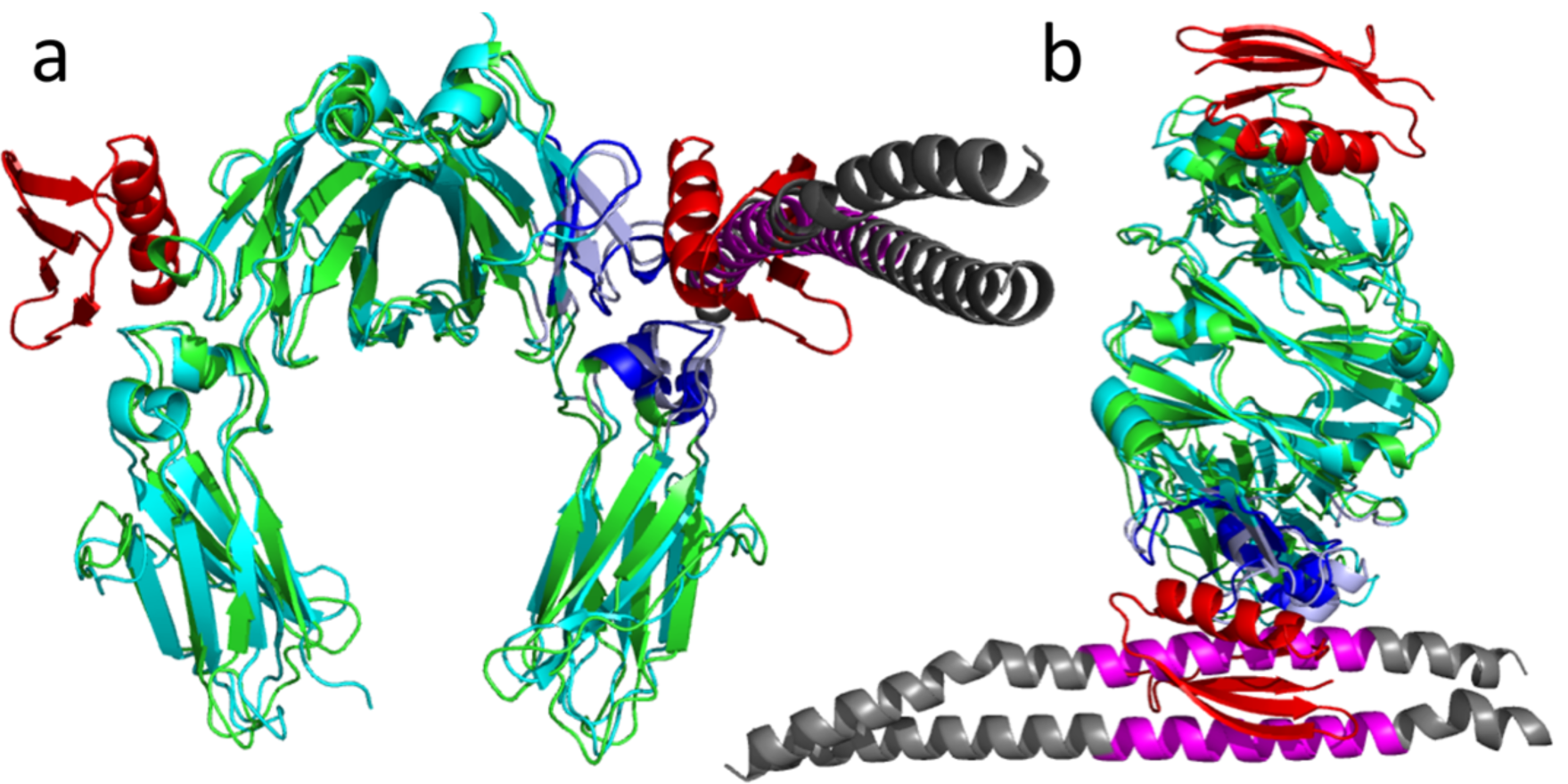
The TX-MS based M1-IgG model aligned with that of the streptococcal protein G. **(a-b)** Protein G (PDB ID: 1FCC) is shown in red with its binding interface on IgG1 in dark blue. The M1 protein is shown in magenta and dark grey with its binding interface on IgG in light blue. Both proteins share the same binding site on IgGs in CH3 domains and close to CH2.

We have additionally made a high-resolution model for the full-length IgG1 interaction with the M1 protein. The model, shown in **Figure 2**, interestingly supports Fab-mediated binding as well, either facilitated by the Fc-binding positioning the Fab-domain in close proximity to the M1 protein leading to the formation of an XL, or by specific, Fab-mediated immune binding. The latter binding model is supported by our previous findings, according to which pooled normal human plasma contains M protein-specific IgG antibodies [4]. Here, we hence extend our earlier results to include the identification of the specific epitopes targeted by these. In addition to the XLs supporting Fc-mediated binding, which is mostly converged to the A- and C1-domains of M1, the immune Fab-interaction is detected on the A- and C1-domains of the M1 protein based on the TX-MS data (**Figure 4**).

**Figure 4:**
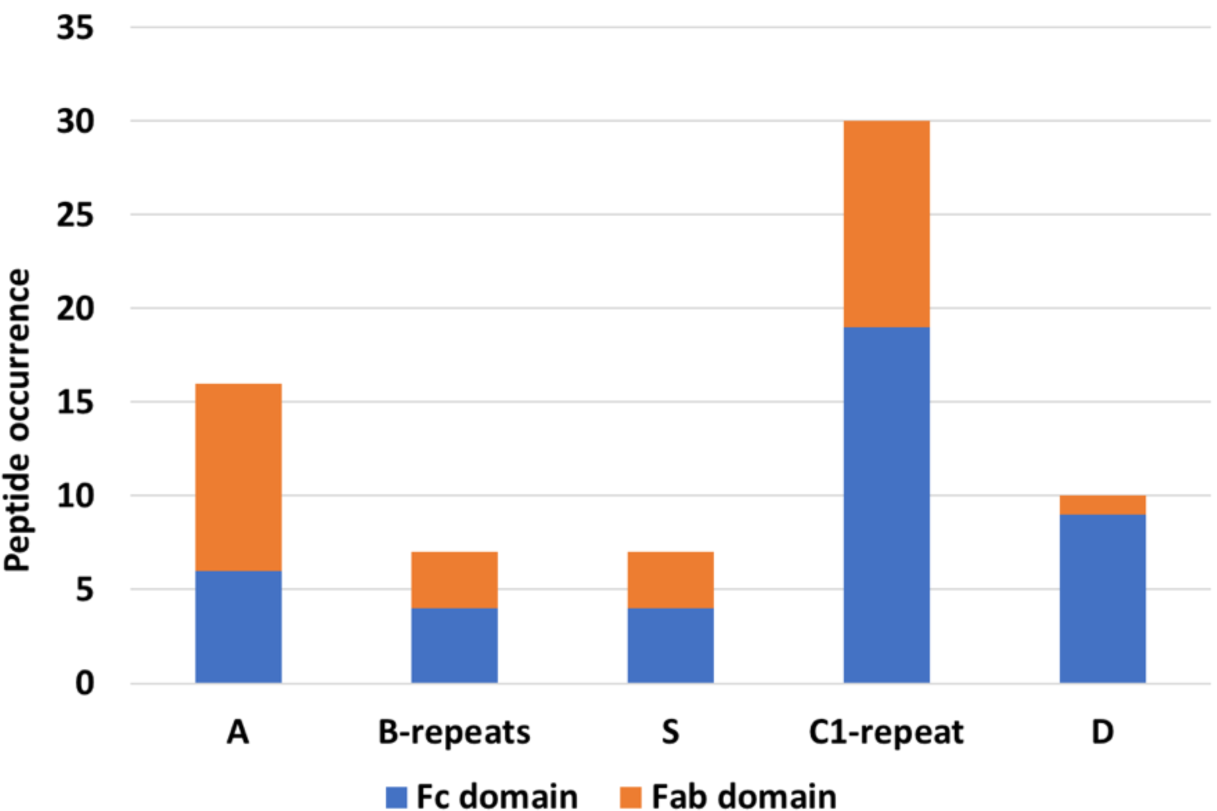
DDA-based peptide quantification of Fc and Fab fragments of IgG subclasses bind to different sub-domains of M1 protein. All IgG subclasses bind to the M1 protein through their Fc-domains. The S- & C1-domains and the hypervariable domain A of the M1 protein have the highest affinity for IgG subclasses. Only IgG heavy chains were considered.

### Comparison of IgG subclasses by XL-MS quantification

Analyzing the DDA data revealed 17 inter-XLs between the M1 protein and the different subclasses of IgGs. However, the distribution of XL-peptides differed between samples with different DSS concentrations. While some of the XLs were detected in all samples (e.g., XLs in **Figure 2**), others were less frequent. We took advantage of this, and quantified the number of detected XL peptides per acquisition specific to each IgG subclass, disregarding that some peptides are identical between all subclasses (***Supplementary Figure 2***). Based on this XL-peptide quantification, two regions of the M1 protein were identified with the highest number of detectable M1-IgG XLs: i) the hypervariable domain A and ii) the S-& C1-domains (end of the S-domain and beginning of the C1-repeat). These important domains of the M1 protein are highly conserved in most types of M proteins [34]. While most of the peptides from the Fc-domain were linked to the S- and C1-domains, VH located Fab-domain peptides were linked to the A-domain as well. **Figure 4** shows the quantification results, where the importance of the C1-repeat and A domain is evident. **Table 1** contains a list of important peptides in the binding interfaces sorted by their occurrences in cross-linking data.

**Table 1:**
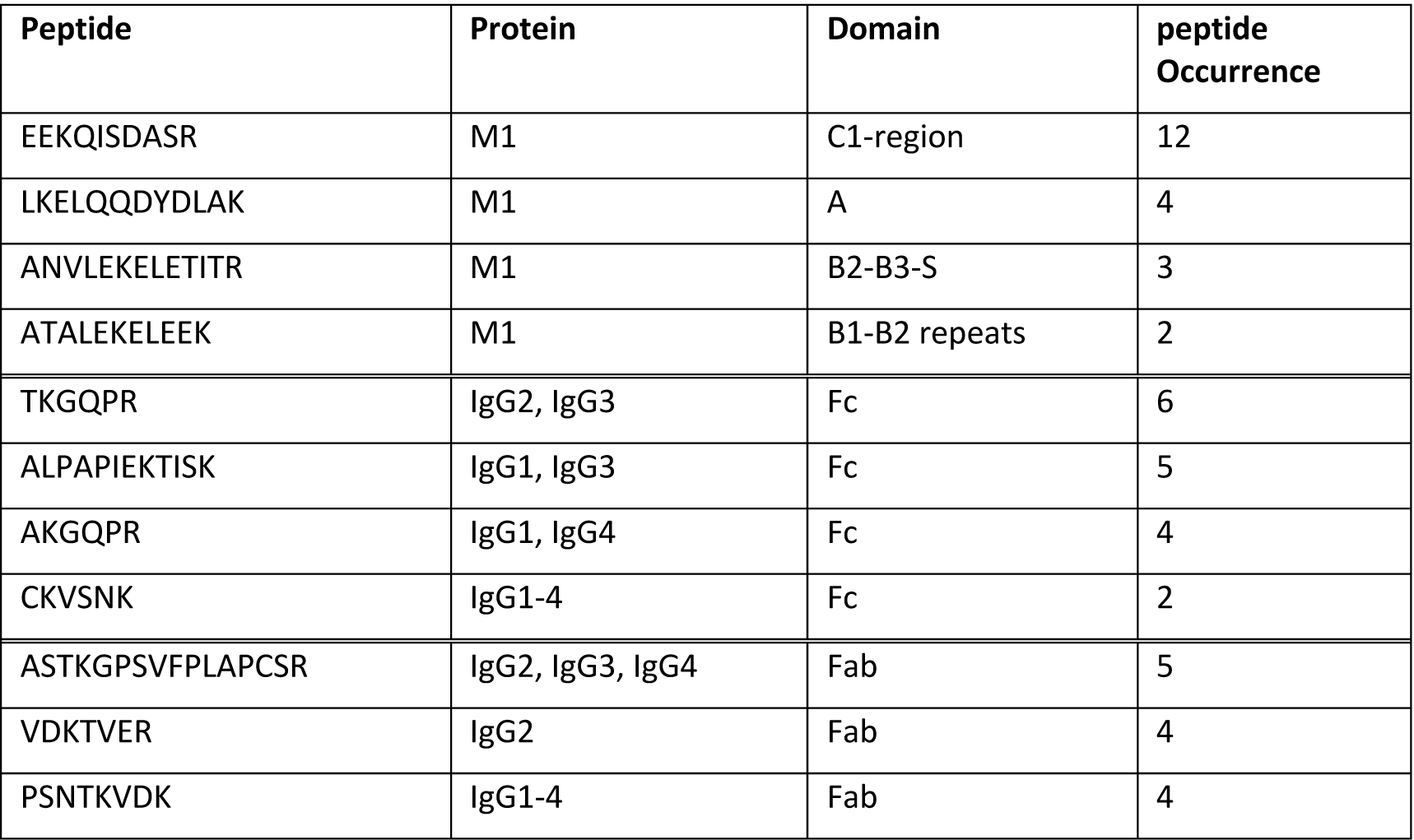
Peptides with high-affinity binding occurrences. The listed peptides are frequent in the binding interfaces of the M1 protein and different IgGs according to the XL-MS quantification.

Interestingly, the detected peptide from the A-domain is homologous to the streptococcal protein G helix, shown to bind IgG1 (**Figure 3** and ***Supplementary Figure 3***). The B-repeats had the fewest number of XLs; however, we identified XLs between B1- and B2-repeats and human plasma fibrinogen in all samples. Fibrinogen is an abundant plasma protein known to bind to the B-repeats of the M proteins [14, 15]. Our results suggest that fibrinogen can mask the IgG epitopes in this region or that very few IgG binds to this region, as indicated in previously published work [35].

### Communication networks

To further evaluate the stability and affinity of the binding interface and to understand the interaction networks between the M1 protein and IgG molecules, MD simulations were carried out. We performed two replicates of 500 ns MD simulations starting from each of the two models for M1(S-domain peptide)-IgG1(Fc) and M1(A-domain peptide)-IgG1(Fc). Based on the simulations, the M1(S)-peptide was stable in one replicate (***Supplementary Movie S1***), while in the other replicate, it completely detached from the Fc-domain after 100 ns (***Supplementary Movie S2***). On the other hand, the M1(A)-peptide remained stable with the Fc-domain during the 500 ns simulation in both replicates (***Supplementary Movie S3*** and ***Supplementary Movie S4***). Next, we performed hydrogen H-bond analysis over the replicates of each system and recorded those that are present for more than 50% of the simulation time at the interface between the IgG1(Fc)- and M1-peptides (**Figure 5-a,b** and ***Supplementary Table 1***). This analysis revealed a strong network of interactions between the M1(A)-peptide-IgG1(Fc), supported by several H-bonds with persistency in the range of 61%-91%. However, we obtained a less stable or transient binding between the M1(S)-peptide-IgG1(Fc) supported by two H-bonds only presented at one replicate.

**Figure 5:**
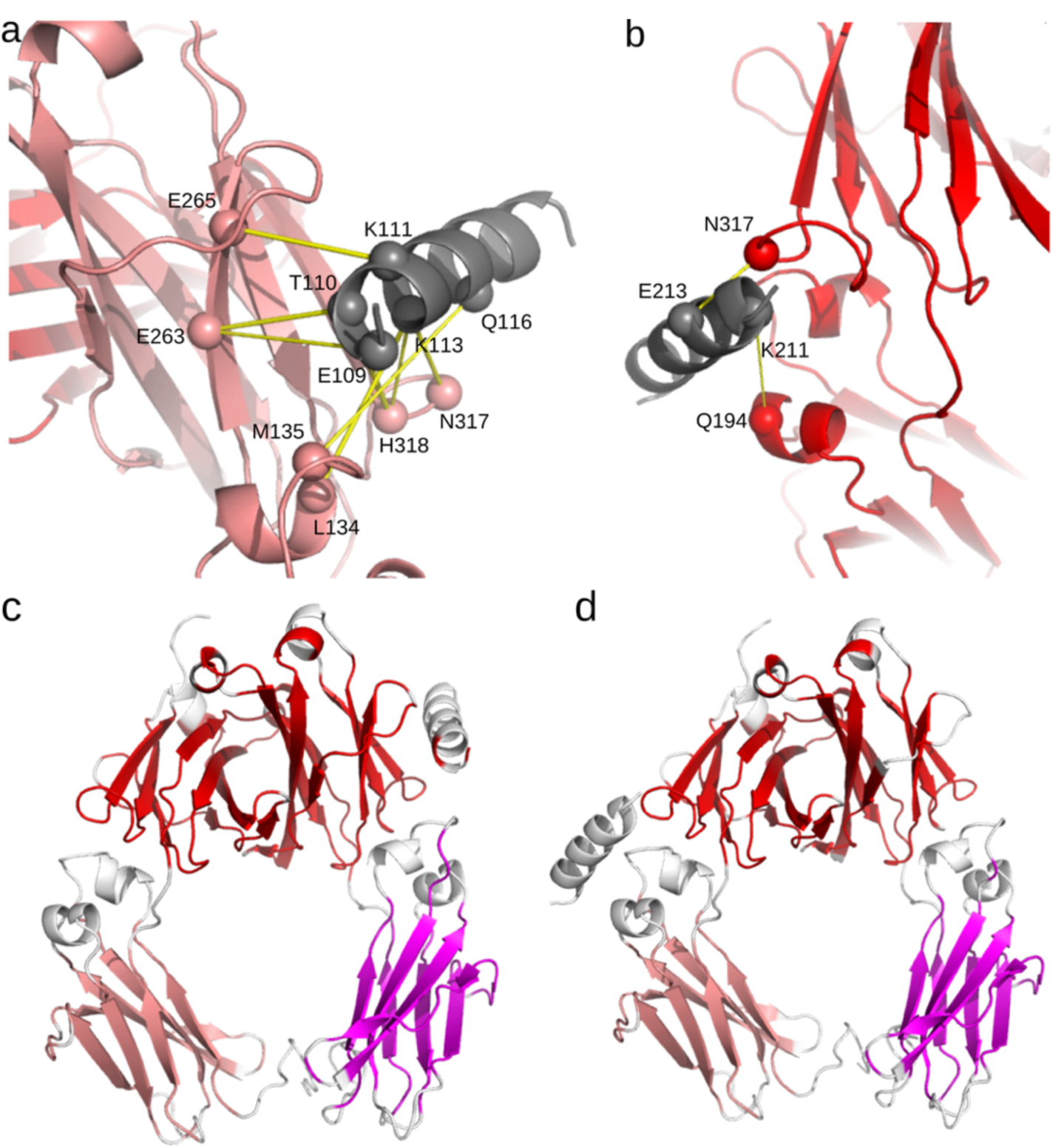
H-bond network between the M1 peptide and IgG Fc-domain. The residues forming H-bond contacts between the **(a)** M1(S)-IgG(Fc) and **(b)** M1(A)-IgG(Fc) are shown with spheres. The thickness of the yellow lines corresponds to the persistence of the interactions throughout the MD simulations. The M1 peptide is colored in gray while the chains of the IgG Fc-domain are depicted in red and pink. **(c and d)** Communication Blocks (CBs^path^) identified by COMMA2 are mapped on the structure of **(c)** M1(S)-IgG(Fc), and **(b)** M1(A)-IgG(Fc). Distinct CBs^path^ of four residues or more are colored in shades of red.

Moreover, for each system, we applied COMMA2 to extract communication blocks, *i.e.*, CBs^path^ (see Methods). The residues comprised in a CB^path^ are linked by communication pathways by transitivity. A communication pathway is defined as a chain of residues displaying correlated motions and linked by stable non-covalent interactions. Hence, it represents an efficient route of information transmission supported by physical interactions. COMMA2 analysis revealed three different CBs^path^ for each system (**Figure 5-c,d**), of which one (in red) contains both CH3 domains of Fc-domain. Two other CBs^path^ are formed (in pink and magenta), one over each CH2 domain. Interestingly, as shown in **Figure 5-c**, two residues from the M1(A)-peptide are in the red block, supporting a strong communication between the M1(A) and IgG1(Fc).

## Discussion

In this study, we characterized the GAS-M1 human-IgG interactions using TX-MS combined with MD simulations. Accordingly, TX-MS revealed the two binding interfaces, while the MD simulations helped to elucidate the interaction network in molecular detail. We identified that the M1 protein is capable of capturing human IgGs to prevent opsonization. All IgG subclasses bind to the M1 protein in a specific region in the CH3 domain and close to CH2, which is involved in binding IgG-receptors (FcγR). This CH2-CH3 binding site has previously been shown to bind streptococcal protein G [31, 32]. Here, the interaction network revealed by MD simulations indicated that both the M1 protein and the streptococcal protein G share the same binding site on the IgG molecules. This interaction would mask the recognition site for FcγR receptors, hence protecting the bacteria from phagocytic killing.

The structural details of the M1 protein in complex with human IgGs revealed important peptides at the binding interface that plays a crucial role in the interactions. We quantified the number of detected XL peptides per sample for each IgG subclass and identified two domains of the M1 protein with the highest affinity to bind IgGs, namely the hypervariable A- and the S-domains (together with the beginning of the C1-repeat). Moreover, our 2 **μ**s MD simulations put in evidence a transient binding for the M1(S)-peptide, and a stable binding for the M1(A)-peptide to the IgG1(Fc)-domain. These results indicated that the two specific peptides of the M1 protein could effectively inhibit the binding of FcγR receptors, making them potential vaccine candidates for future studies.

Most vaccine developments and trials targeted for *S. pyogenes* are currently focused on the M proteins and their fragments, with the leading ones targeting the N-terminal or the C-repeat region [36]. We have previously shown that peptides in the M protein C-region containing the EEKQISDASR- motif (**Table 1**) are potent epitopes for opsonizing antibodies and crucial for the interaction and internalization with monocytic cells; a result corroborated here [4]. Our data also indicates that epitopes in the A-domain are central in both Fab- and Fc-IgG1-mediated evasion of the human immune responses, and potential novel targets for therapeutic strategies to combat GAS infections.

## Conclusions

GAS is an important human pathogen infecting more than 700 million individuals globally each year. Here, we describe the interaction of the M1 protein, an important virulence factor of GAS, with human immunoglobulin G (IgG1-4) using targeted cross-linking mass spectrometry combined with molecular dynamic simulations. These interactions revealed that all IgG subclasses could bind to the M1 protein with their non-immune Fc-domains and share roughly the same Fc-binding interface on the Fc-receptor-binding domain, showing the crucial role of the M1 protein to eliminate IgG-Fc receptors (FcγR) interactions and protect the bacteria from phagocytic killing. Finally and as the result of this study, the highly frequent peptides in the important C1-repeat (EEKQISDASR), the transient binding peptide of S-domain (GNAKLELDQLSSEKEQ), and the strong binding peptide in the hypervariable A-domain (LETKLKELQQDYDLAK) on the interface of the M1 protein and IgGs can be used as vaccine candidates for further studies.

## Acknowledgment

This work was supported by Foundation of Knut and Alice Wallenberg (2016.0023) to LH, JM and LM; Swiss National Science Foundation (grant no. P2ZHP3_191289) to HK and GTVN; The Fondation pour la Recherche Médicale (Equipe FRM 2017M.DEQ20170839114) to YK and MN.

## Author Contributions

HK, LH, JM, and LM conceived the study. HK and LH wrote the manuscript. HK generated computational models, analyzed and interpreted MS data and wrote the first draft. LH designed and performed cross-linking MS sample preparation and analysis. YK designed and performed MD analysis and interpreted the data and approved the final draft. LM, JM, GTVN, and MN interpreted data, contributed to the manuscript preparation and approved the final draft; All of the authors have read and approved the final manuscript.

## Supplementary Materials

**Supplementary Table 1:**
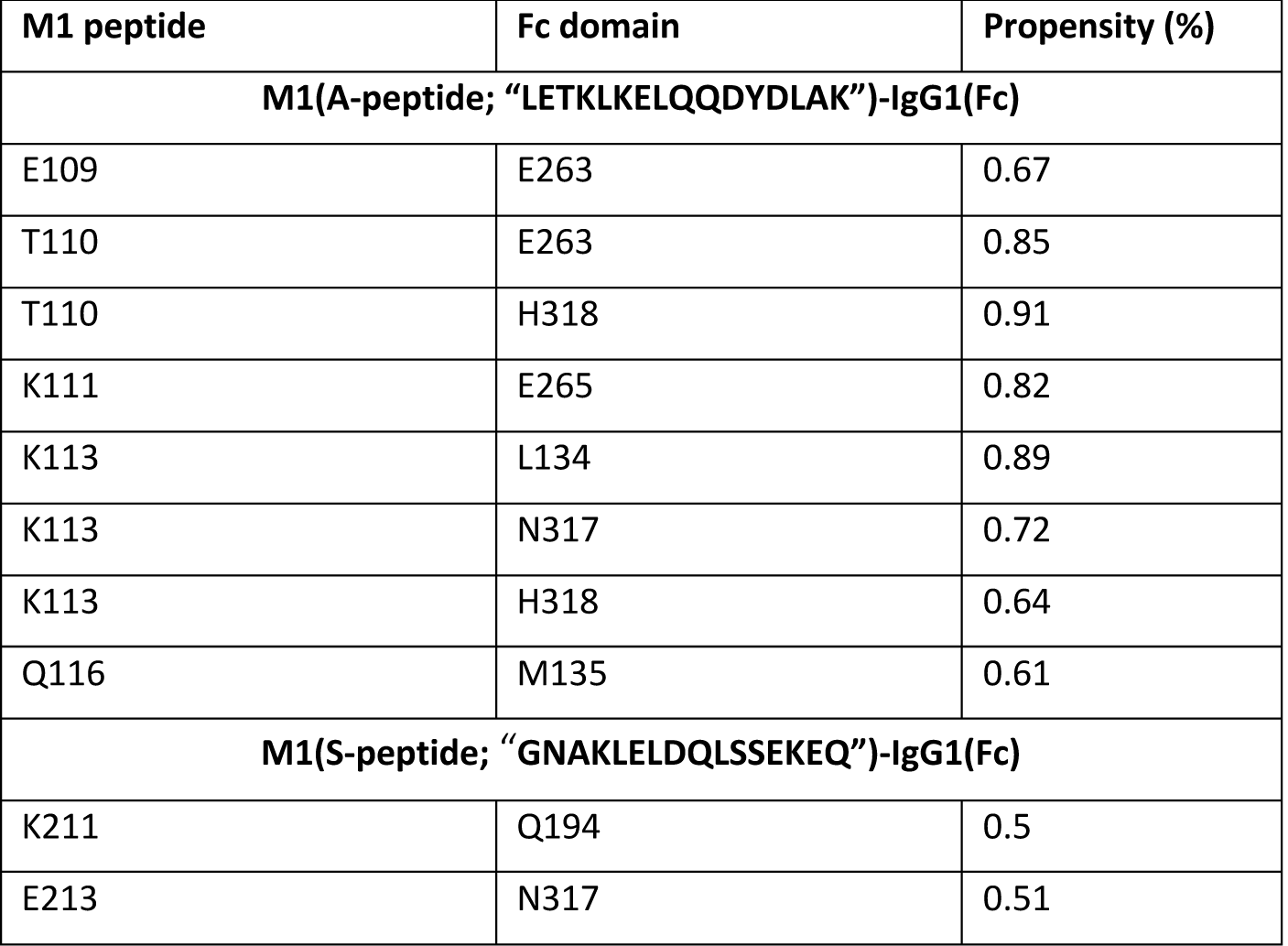
H-bonds formed between the M1 peptides and IgG1 Fc domain. The persistence of the H-bonds formed at the interface are recorded as a percentage over the MD simulation time. For each studied system, the maximum values are reported over the two replicates.

**Supplementary Figure 1:**
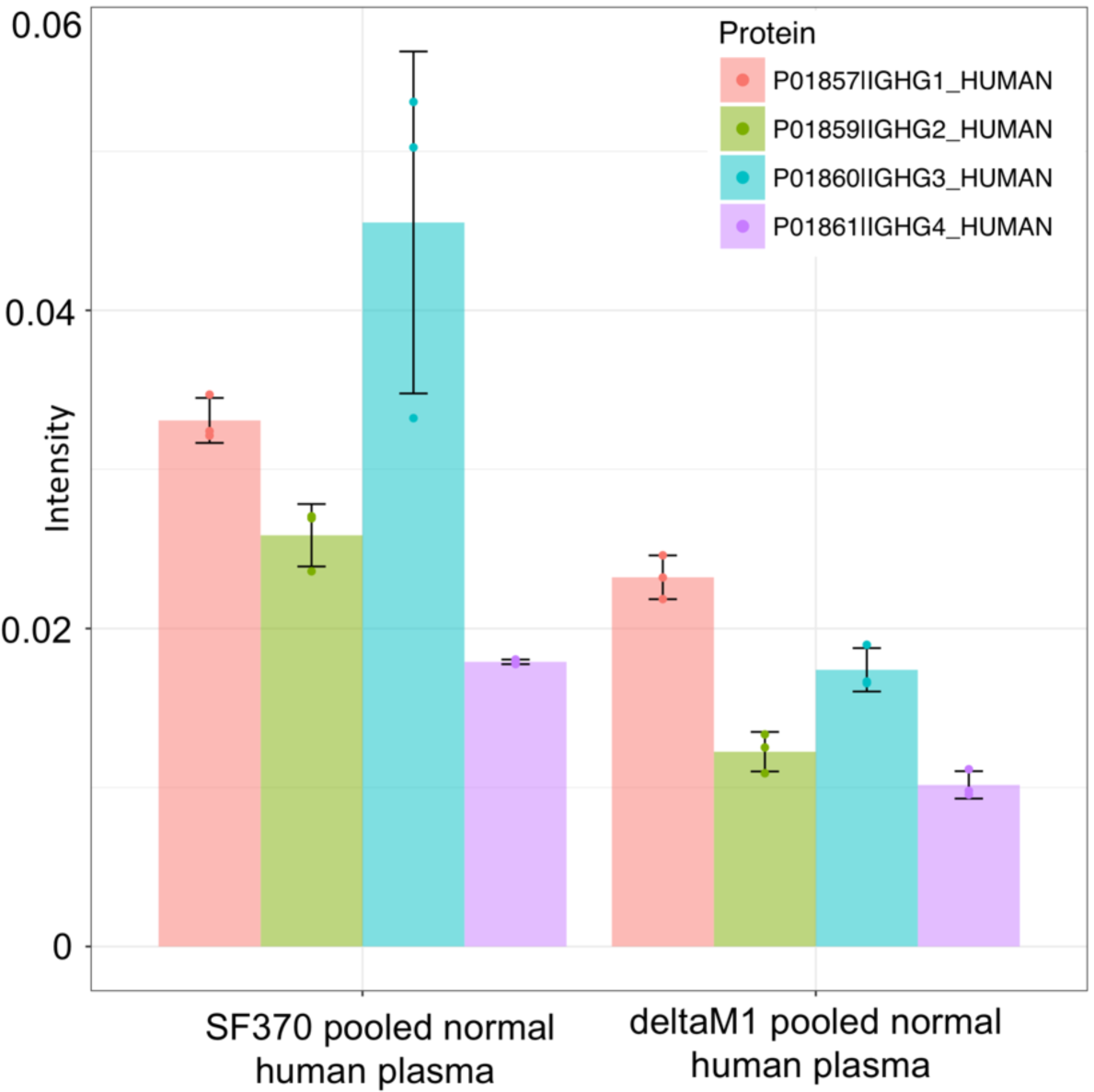
TIC normalized intensity of human IgG subclasses in plasma samples. The intensity level of the heavy chains of human IgGs is analyzed through a DIA-MS analysis approach. Two groups of samples are considered here: pooled normal human plasma on the surface of the wt strain SF370 and on the surface of an SF370-derived M1 mutant strain (deltaM1). The heavy chains of IgG3 and IgG4 (IGHG3 and IGHG4, respectively) have the highest and the lowest intensities among all IgG subclasses. The data also indicates that *S. pyogenes* has a high affinity for IgG1 and IgG3, which is M1-mediated.

**Supplementary Figure 2:**
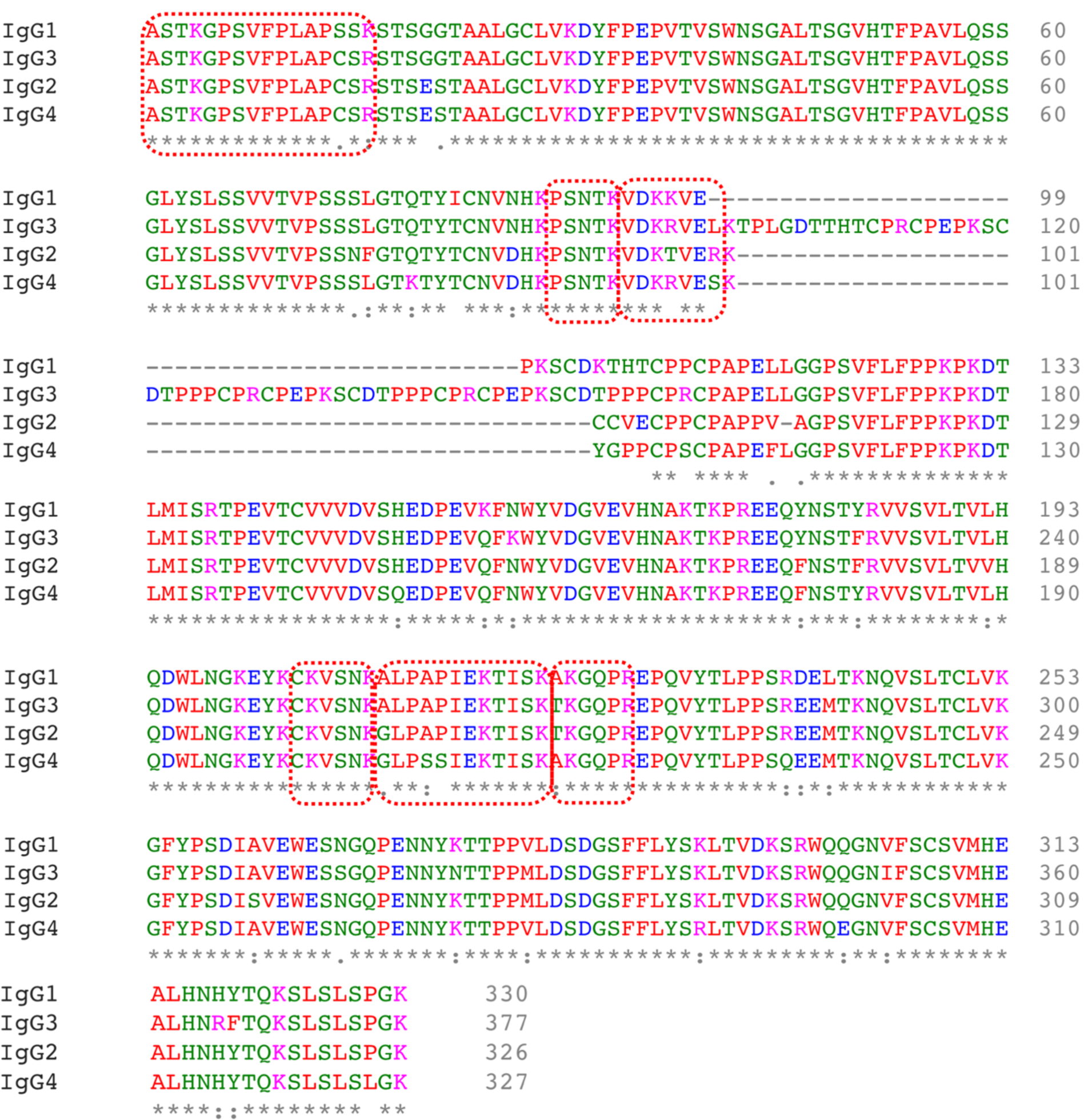
Multiple sequence alignment of all IgG subclasses. Identified XLs are shown with red dashed boxes where small sequence differences can be seen. The main difference can be noticed between IgG3 and other subclasses as the longer hinge region of IgG3 (residues 100 - 150 in IgG3).

**Supplementary Figure 3:**
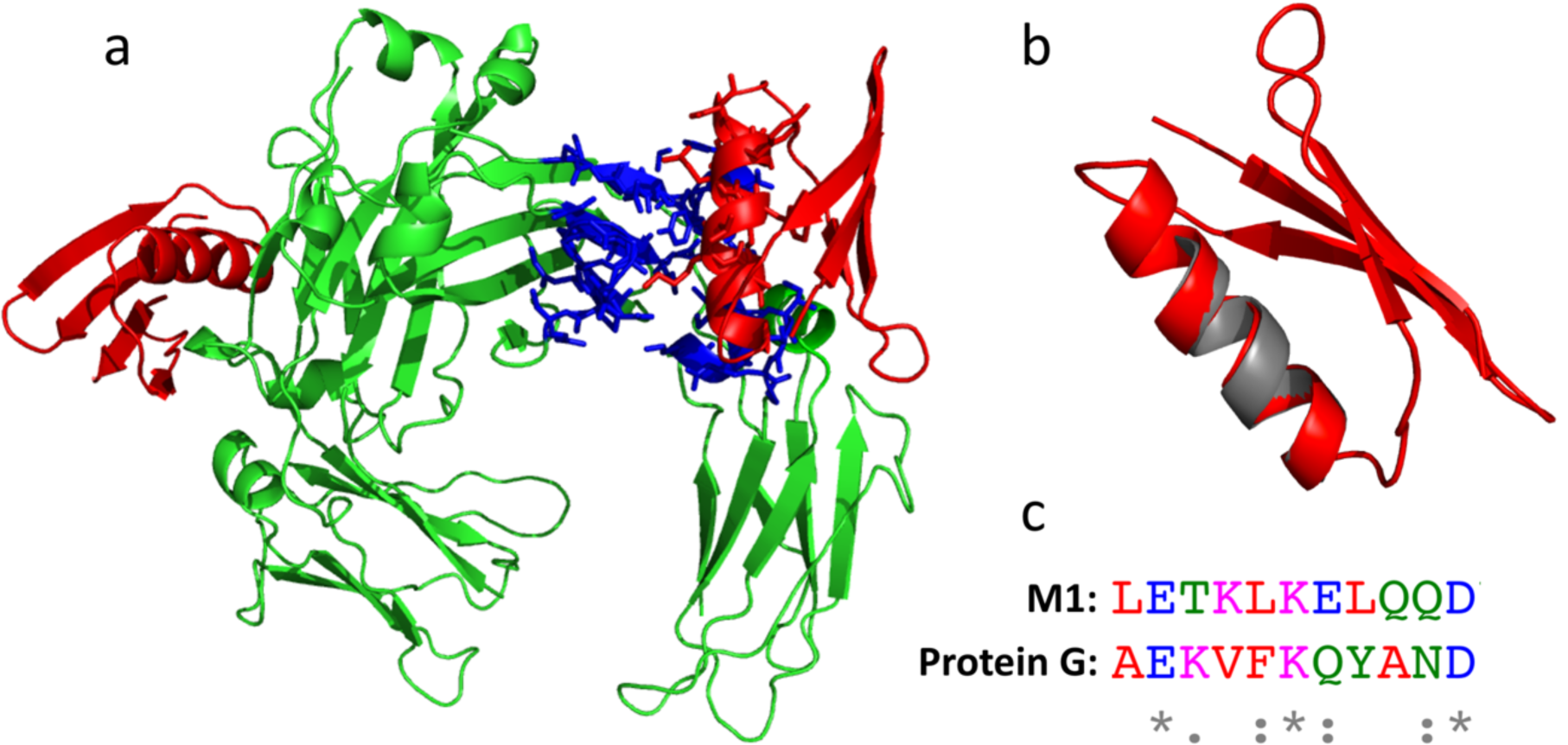
Sequence similarity between protein G helix and the detected peptide of M1-A domain. **(a)** Crystal structure of protein G and human IgG1 (PDB id 1fcc). **(b, c)** Structural and sequence alignment of protein G helix (in red) on the peptide from the M1-A domain (in grey) detected by cross-linking mass spectrometry as the high-affinity peptide to bind IgGs.

**Supplementary Figure 4:**
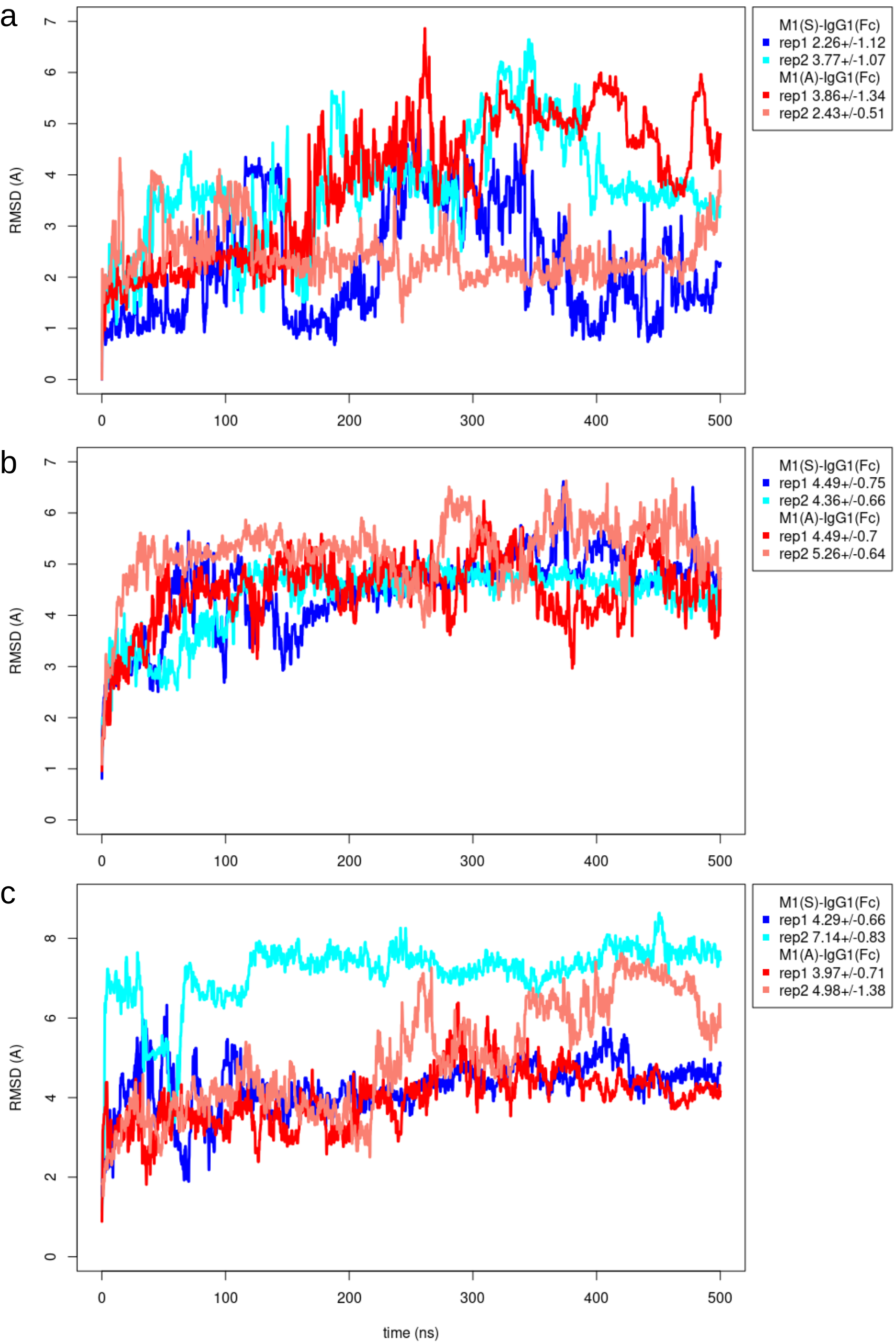
The root mean square deviations for M1(S)-IgG(Fc) and M1(A)-IgG(Fc). The RMSD from the equilibrated structure is computed on the backbone (C, Ca, N, O) atoms for the M1 peptides **(a)** and the two chains of Fc **(b** and **c)**. The two replicates of M1(S)-IgG(Fc) are colored in blue and cyan, and the replicates of M1(A)-IgG(Fc) in red and pink.

**Supplementary Figure 5:**
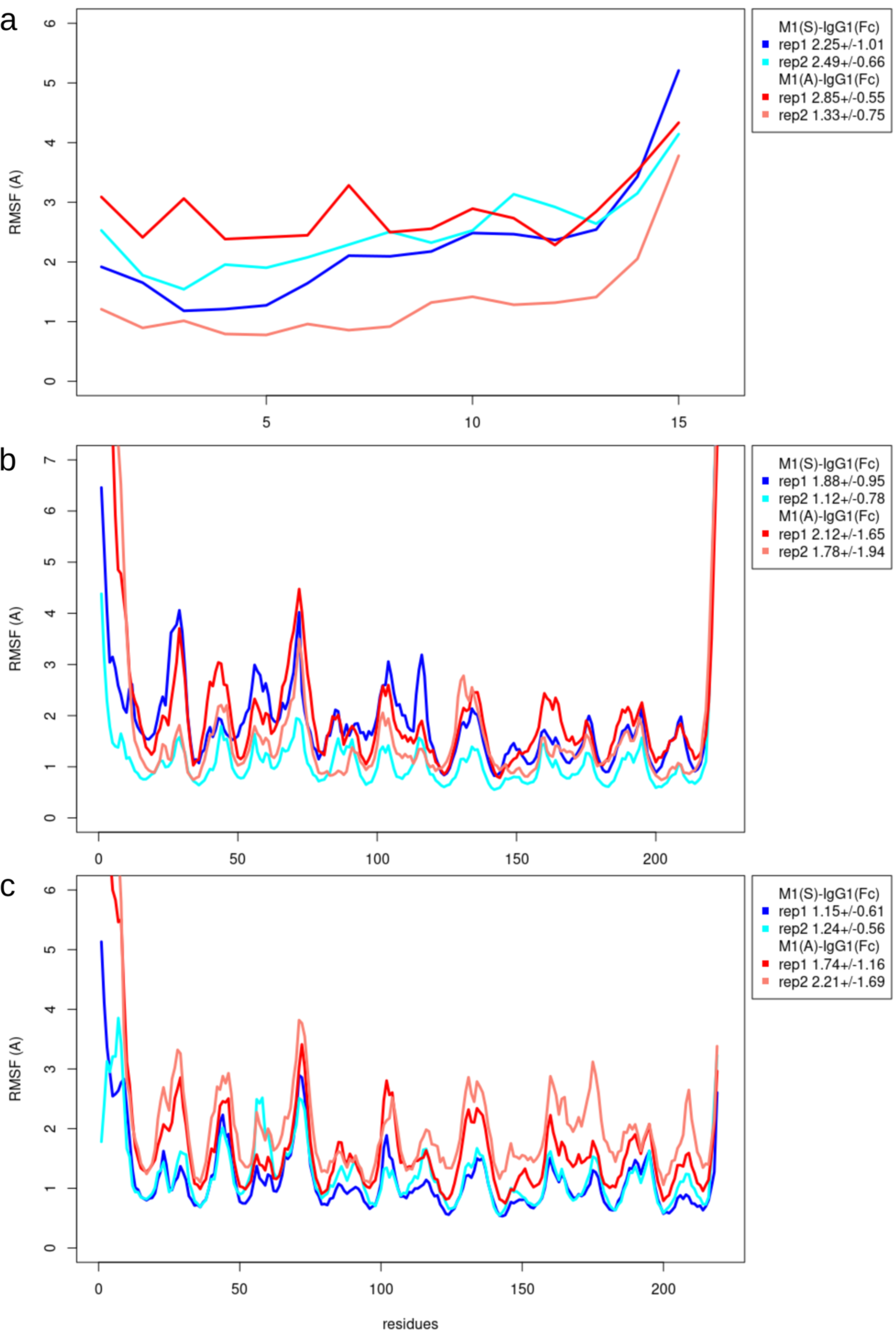
The root mean square fluctuations for M1(S)-IgG(Fc) and M1(A)-IgG(Fc). The RMSF was measured on the backbone (C, Ca, N, O) atoms and averaged by residue, considering the last 400 ns of the MD simulations for the M1 peptides **(a)** and the two symmetrical chains of Fc **(b** and **c)**. The two replicates of M1(S)-IgG(Fc) are colored in blue and cyan, and the replicates of M1(A)-IgG(Fc) in red and pink.

**Supplementary Movie S1: 500 ns MD simulation of M1(S)-IgG1(Fc), first replicate.**

**Supplementary Movie S2: 500 ns MD simulation of M1(S)-IgG1(Fc), second replicate.**

**Supplementary Movie S3: 500 ns MD simulation of M1(A)-IgG1(Fc), first replicate.**

**Supplementary Movie S4: 500 ns MD simulation of M1(A)-IgG1(Fc), second replicate.**

## References

[1] T. Mitchell, “The pathogenesis of streptococcal infections: from Tooth decay to meningitis.,” Nat Rev Microbiol, vol. 1, pp. 219–230, 2003.

[2] A. Ralph and J. Carapetis, “Group a streptococcal diseases and their global burden.,” Curr Top Microbiol Immunol., vol. 368, p. 1–27, 2013.

[3] M. Wisniewska, L. Happonen, F. Kahn, M. Varjosalo, L. Malmström, G. Rosenberger, C. Karlsson, G. Cazzamali, I. Pozdnyakova, I. Frick, L. Björck, W. Streicher, J. Malmström and M. Wikström, “Functional and structural properties of a novel protein and virulence factor (Protein sHIP) in Streptococcus pyogenes.,” J Biol Chem., vol. 289, no. 26, p. 18175–18188, 2014.

[4] L. Happonen, S. Hauri, G. S. Birkedal, C. Karlsson, T. d. Neergaard, H. Khakzad, P. Nordenfelt, M. Wikström, M. Wisniewska, L. Björck, L. Malmström and J. Malmström, “A quantitative Streptococcus pyogenes–human protein–protein interaction map reveals localization of opsonizing antibodies,” Nature Communications, vol. 10, no. 2727, 2019.

[5] S. Chowdhury, L. Happonen, H. Khakzad, L. Malmström and J. Malmström, “Structural proteomics, electron cryo-microscopy and structural modeling approaches in bacteria–human protein interactions.,” Medical Microbiology and Immunology, 2020.

[6] U. Pawel-Rammingen von, B. P. Johansson and L. Björck, “IdeS. a novel streptococcal cysteine proteinase with unique specificity for immunoglobulin G.,” EMBO J., vol. 21, pp. 1607–1615, 2002.

[7] M. Collin and A. Olsén, “EndoS, a novel secreted protein from Streptococcus pyogenes with endoglycosidase activity on human IgG.,” EMBO J., vol. 20, p. 3046–3055, 2001.

[8] S. J., W. B. Struwe, E. Cosgrave, P. Rudd, M. Stervander, M. Allhorn, A. Hollands, V. Nizet and M. Collin, “EndoS2 is a unique and conserved enzyme of serotype M49 group A Streptococcus that hydrolyses N-linked glycans on IgG and α1-acid glycoprotein.,” Biochem. J., vol. 455, pp. 107–118, 2013.

[9] L. Björk and G. Kronvall, “Purification and Some Properties of Streptococcal Protein G, a Novel IgG-binding Reagent,” J Immunol., vol. 133, no. 2, pp. 969–974, 1984.

[10] K. J. Reis, E. M. Ayoub and M. D. Boyle, “Streptococcal Fc Receptors. I. Isolation and Partial Characterization of the Receptor From a Group C Streptococcus,” J Immunol., vol. 132, no. 6, pp. 3091–3097, 1984.

[11] D. G. Heath and P. P. Cleary, “Fc-receptor and M-protein Genes of Group A Streptococci Are Products of Gene Duplication,” Proc. Natl. Acad. Sci., vol. 86, no. 12, pp. 4741–4745, 1989.

[12] G. Vidarsson, G. Dekkers and T. Rispens, “IgG Subclasses and Allotypes: From Structure to Effector Functions,” Frontiers in Immunology, vol. 5, p. 520, 2014.

[13] K. Sjöholm, C. Karlsson, A. Linder and J. Malmström, “A comprehensive analysis of the Streptococcus pyogenes and human plasma protein interaction network,” Mol. Biosyst., vol. 10, p. 1698–1708, 2014.

[14] P. Macheboeuf, C. Buffalo, C. Fu, A. Zinkernagel, J. Cole, J. Johnson, V. Nizet and P. Ghosh, “Streptococcal M1 protein constructs a pathological host fibrinogen network.,” Nature, vol. 472, no. 7341, pp. 64–68, 2011.

[15] S. Hauri, H. Khakzad, L. Happonen, J. Teleman, J. Malmström and L. Malmström, “Rapid determination of quaternary protein structures in complex biological samples,” Nature Communications, vol. 10, no. 192, 2019.

[16] P. Nordenfelt, S. Waldemarson, A. Linder, M. Mörgelin, C. Karlsson, J. Malmström and L. Björck, “Antibody orientation at bacterial surfaces is related to invasive infection.,” The Journal of experimental medicine, vol. 209, no. 13, p. 2367–2381, 2012.

[17] J. Koehler Leman, B. Weitzner, S. Lewis and a. et, “Macromolecular Modeling and Design in Rosetta: New Methods and Frameworks.,” Nature Methods, 2020.

[18] Y. Karami, F. Guyon, S. De Vries and P. Tufféry, “DaReUS-Loop: accurate loop modeling using fragments from remote or unrelated proteins,” Scientific Reports, vol. 8, no. 1, p. 13673, 2018.

[19] Y. Karami, J. Rey, G. Postic, S. Murail, P. Tufféry and S. J. de Vries, “DaReUS-Loop: a web server to model multiple loops in homology models,” Nucleic Acids Research, vol. 47, no. W1, pp. W423–W428, 2019.

[20] J. Söding, A. Biegert and A. Lupas, “The HHpred interactive server for protein homology detection and structure prediction.,” Nucleic Acids Res., vol. 33(Web Server issue), p. W244–W248, 2005.

[21] Y. Song, F. DiMaio, R. Wang, D. Kim, C. Miles, T. Brunette, J. Thompson and D. Baker, “High-resolution comparative modeling with RosettaCM.,” Structure., vol. 21, no. 10, pp. 1735–1742, 2013.

[22] R. F. Alford, A. Leaver-Fay, J. R. Jeliazkov, M. J. O’Meara, F. P. DiMaio, H. Park, M. V. Shapovalov, P. D. Renfrew, V. K. Mulligan, K. Kappel, J. W. Labonte, M. S. Pacella and R. Bonneau, “The Rosetta All-Atom Energy Function for Macromolecular Modeling and Design,” Journal of Chemical Theory and Computation, vol. 13, no. 6, pp. 3031–3048, 2017.

[23] K. Sjöholm, O. Kilsgård, J. Teleman, L. Happonen, L. Malmström and J. Malmström, “Targeted Proteomics and Absolute Protein Quantification for the Construction of a Stoichiometric Host-Pathogen Surface Density Model,” American Society for Biochemistry and Molecular Biology, vol. 16, no. 4 suppl 1, pp. S29–S41, 2017.

[24] J. J. Gray, “High-resolution protein–protein docking,” Current Opinion in Structural Biology, vol. 16, no. 2, pp. 183–193, 2006.

[25] D. Van Der Spoel, E. Lindahl, B. Hess, G. Groenhof, A. E. Mark and H. J. Berendsen, “GROMACS: fast, flexible, and free,” J Comput Chem, vol. 26, no. 16, pp. 1701–1718, 2005.

[26] M. A. Martí-Renom, A. C. Stuart, A. Fiser, R. Sánchez, F. Melo and A. Sali, “Comparative protein structure modeling of genes and genomes,” Annu Rev Biophys Biomol Struct, vol. 29, pp. 291–325, 2000.

[27] R. J. Loncharich, B. R. Brooks and R. W. Pastor, “Langevin dynamics of peptides: The frictional dependence of isomerization rates of N-acetylalanyl-N’-methylamide,” Biopolymers, vol. 32, no. 5, pp. 523–535, 1992.

[28] T. Darden, D. York and L. Pedersen, “Particle mesh Ewald: An Nlog(N) method for Ewald sums in large systems,” The Journal of Chemical Physics, vol. 98, pp. 10089–10092, 1993.

[29] Y. Karami, E. Laine and A. Carbone, “Dissecting protein architecture with communication blocks and communicating segment pairs,” BMC bioinformatics, vol. 17, no. 2, 2016.

[30] Y. Karami, T. Bitard-Feildel, E. Laine and A. Carbone, “Infostery” analysis of short molecular dynamics simulations identifies highly sensitive residues and predicts deleterious mutations,” Scientific reports, vol. 8, no. 1, pp. 1–18, 2018.

[31] A. Sauer-Eriksson, G. Kleywegt, M. Uhlén and T. Jones, “Crystal structure of the C2 fragment of streptococcal protein G in complex with the Fc domain of human IgG.,” Structure, vol. 3, p. 265–278, 1995.

[32] B. Akerström, T. Brodin, K. Reis and L. Björck, “Protein G: a powerful tool for binding and detection of monoclonal and polyclonal antibodies.,” The Journal of Immunology, vol. 135, no. 4, p. 2589, 1985.

[33] J. Deisenhofer, “Crystallographic refinement and atomic models of a human Fc fragment and its complex with fragment B of protein A from Staphylococcus aureus at 2.9- and 2.8-Å resolution.,” Biochemistry, vol. 20, p. 2361–2370, 1981.

[34] H. Khakzad, J. Malmström and L. Malmström, “Greedy de novo motif discovery to construct motif repositories for bacterial proteomes,” BMC Bioinformatics, vol. 20, no. 141, 2019.

[35] J. Lannergård, B. M. Kristensen, M. C. U. Gustafsson, J. J. Persson, A. Norrby-Teglund, M. Stålhammar-Carlemalm and G. Lindahl, “Sequence variability is correlated with weak immunogenicity in Streptococcus pyogenes M protein,” MicrobiologyOpen, vol. 4, no. 5, pp. 774–789, 2015.

[36] V. Ozberk, M. Pandey and M. F. Good, “Contribution of Cryptic Epitopes in Designing a Group A Streptococcal Vaccine,” Hum Vaccin Immunother., vol. 14, no. 8, pp. 2034–2052, 2018.

[37] B. D. Weitzner, J. R. Jeliazkov, S. Lyskov, N. Marze, D. Kuroda, R. Frick, J. Adolf-Bryfogle, N. Biswas, R. L. J. Dunbrack and J. J. Gray, “Modeling and docking of antibody structures with Rosetta,” Nature protocols, vol. 12, no. 2, pp. 401–416, 2017.

[38] K. Wenig, L. Chatwell, U. von Pawel-Rammingen, L. Björck, R. Huber and P. Sondermann, “Structure of the streptococcal endopeptidase IdeS, a cysteine proteinase with strict specificity for IgG.,” Proceedings of the National Academy of Sciences, vol. 101, no. 50, pp. 17371–17376, 2004.

